# The effect of nutrient storage on courtship behavior and copulation frequency in the fruit fly, *Drosophila melanogaster*

**DOI:** 10.1101/331660

**Authors:** Hannah Ananda Bougleux Gomes, Justin R. DiAngelo, Nicholas Santangelo

## Abstract

Nutrient storage and metabolism effects on reproductive behavior are well studied in higher vertebrates like mammals, but are less understood in simpler systems. *Drosophila melanogaster* is well suited to study the ramifications of diet and metabolic energy storage on reproductive behaviors as they are commonly used to explore energy mobilization pathways. We tested, for the first time, courtship of the naturally occurring *adipose (adp^60^)* mutant which over-accumulates triglycerides and glycogen on a normal diet. We also fed wild type (WT) flies either a normal diet, high fat diet or food deprived them before measuring courtship, copulations, and glycogen and triglyceride levels. *Adipose* mutants decreased both courtship and copulation frequency, yet showed the highest glycogen and triglyceride levels. We suggest the *adp^60^* physique and/or an altered ability to utilize mobilize energy explains these effects. Food deprived WT flies had the lowest glycogen and triglycerides but exhibited shortened courtship latencies with increased courtship behaviors. This may be due to a decreased lifespan of food deprived flies leading to a greater reproductive drive. However, high fat fed flies copulated more frequently and had the highest triglycerides among WT groups, yet equal glycogen levels to the normal fed WT group. Thus, a high fat diet either increases male attractivity or male courtship persistence. Taken together, available diet and nutrient storage affects male fly reproductive behavior in a unique manner, which may be explained by their natural history, and provides a paradigm for understanding energetics based on reproductive potential.

## Introduction

Caloric intake is known to play an important role in different organisms’ behaviors. For example, consumption of a high-calorie diet alters the function of the mammalian circadian clock, thereby affecting behavioral processes, such as locomotor activity, sleep and energy homeostasis [1]. In the fruit fly *Drosophila melanogaster*, the presence of food promotes aggressive behaviors in males mediated by sweet-sensing gustatory receptor neurons [2]. Conversely, caloric restriction can also affect behaviors. A long-term experiment with humans in free-living conditions showed that caloric restriction resulted in decreased physical activity levels [3]. Studies such as these show that altering caloric intake not only impacts behavior via activity level decisions, but social interactions as well.

In addition to caloric intake influencing behavior, the storage of energy as glycogen and fat and the ability to mobilize these stores also influences reproductive behaviors. For example, available energy resources during mating of tetrapods are well known to be related to the physiological ability of these animals to carry out reproduction (mammals: reviewed in [4], [5], [6], [7]; birds: reviewed [8], [9]; reptiles: [10], [11]). The clear link between reproductive physiology and energy reserves suggests that appetitive reproductive behaviors should also be tightly linked to these energy reserves. Such a mechanism would enable organisms to adequately assess whether the proper energetic resources are available to complete the process of reproduction once started. This is well known to happen in mammals, where the hormonal pathways that control energy balance are directly linked to the ones that control sexual reproduction (reviewed in [6]). However outside of mammals, data on the link between reproductive behavior and caloric intake and mobilization is lacking (but see [12] and review within on birds). In addition, males are rarely the subject of such studies, yet one would expect that, despite reproduction requiring less energetic output for males, the same selective pressure of monitoring metabolic energy resources in making reproductive decisions should remain. Cheng et al. [13] showed that male oriental fruit flies (*Bactrocera dorsalis*) reared on food with high levels of D-glucose exhibit higher success in mating. Also, Mediterranean fruit flies (C*eratitis capitata*) caught while undergoing reproductive behaviors had higher lipid levels than flies at rest [14] with males having much more variable levels of lipids than females. Yet the effect of energy availability on appetitive reproductive behaviors such as courtship is not as well known. Such data could provide the beginnings of a framework to understand how metabolic energy resources shape appetitive behaviors.

In order to explore any correlation that may exist between reproductive behavior and metabolism, energy storage can be manipulated genetically and/or through diet. The fruit fly *Drosophila melanogaster* has been a useful model system for these types of studies because of its high genetic conservation to humans, its ease of obtaining large sample sizes, the ease of genetic and dietary manipulations, and well-characterized and robust reproductive behaviors that can be easily quantitated. In *D. melanogaster* there exist mutants in metabolic genes that result in lean or obese phenotypes similar to that seen when wild type flies are food deprived or fed a high-fat diet, respectively [15].

These mutants can be utilized to test whether altering metabolic pathways affects behavior. One such gene that can be useful in this regard is *adipose (adp)*. The most well-characterized mutation of this gene is *adp^60^*, a 23 nucleotide deletion resulting in increased storage of triglycerides in the fly fat body, providing an obesity phenotype, similar to the triglyceride accumulation observed when flies are fed a high fat diet [16], [17]. While both the *adp^60^* mutation and feeding flies a high fat diet leads to an obesity phenotype, the *adp^60^* mutation results in chronic obesity as the fat accumulation is observed in both the larval and adult stages of development [16], [18]. This is different than the obesity observed after feeding flies a high fat diet as this obesity is acute and only appears in adult flies after 4-5 days of being exposed to the altered diet [17]. While much is known about the metabolic consequences due to the loss of the *adipose* gene, little is known about how the obesity phenotype resulting from this *adp^60^* mutation affects reproductive behaviors.

Metabolic mutants can potentially have similar changes in metabolism as those fed different diets without manipulating the feeding regimen, thus strengthening the link between energy storage and reproductive behaviors. Both approaches, and reproductive behavior monitoring, can be easily carried out in *D. melanogaster* making them an ideal model for which to explore this paradigm. For example, drive to reproduce can be assessed through courtship behaviors such as wing vibrations, wing scissoring, tapping, thrusting, and copulation attempts and copulations [19]. In this study, we compare these reproductive behaviors from *adp^60^* male flies, wild type males fed a high-fat diet, or wild type males food deprived for 24 hours to wild type male flies fed a normal diet in order to determine the effects of changes in nutrient storage and metabolism on reproductive behaviors. For the purposes of this study, female flies are kept constant (i.e. are only of WT genotype and not diet manipulated) in order to isolate any potential effects of nutrient storage on male courtship decisions.

Therefore, in addressing the effects of obese mutants and high fat feeding and starvation on reproduction, our study is an opportunity to increase the base of knowledge on the behavioral effects of obesity by utilizing the *adp^60^* mutant in addition to the diet-induced obesity model. By comparing the courtship behaviors and frequency of copulation of male flies fed a high-fat diet, males food deprived for 24 hours, and male *adp^60^* mutants, with wild type *D. melanogaster* males fed a normal diet, we aim to understand how energy resources and nutrient availability affect reproductive behaviors. The use of both diet and *adp^60^* mutants will help establish whether any resulting changes in male courtship due to diet regime is related to changes in nutrient storage (i.e. presence or absence of *adp^60^* gene), or is the simply the cue of caloric intake (i.e. high or low caloric diet).

Relative to normal fed wild type flies, we predict *adp^60^* mutant flies, and flies fed a high-fat diet, to show an increase in copulation frequency and an increase in courtship behaviors because of the enhanced energy reserves that could be invested in courtship. This prediction is consistent with data from Kauffmann and Rissman [20] where the hormone GnRH-II mediates sexual behavior in mice when there are enough energy resources present. This helps ensure enough energy is available to mediate successful reproduction. Though invertebrates such as insects may utilize a different hormonal mechanism, we predict the relationship between energetic requirements and successful reproduction to be a relatively conserved relationship. Conversely, we expect food deprived flies to show a decrease in copulation and courtship for similar reasons. We expect that the *adp^60^* mutants show a similar duration of courtship behaviors and courtship latency to the wild type flies fed a high-fat diet and thus that *adp^60^* mutants are in fact a good model for understanding the effects of obesity on outward male sexual behavior.

## Materials and Methods

### Fly husbandry

Flies used in this study were: OreR (BL#2376) and *adp^60^* outcrossed into the OreR genetic background (a gift from Ronald Kuhnlein). For the courtship assays, 1-2 day old male flies were separated from females and aged 4-5 days before being put into the courtship apparatus with 3-5-day old wild type OreR virgin females. For high fat diet studies, OreR males were fed cornmeal-sucrose medium (9g *Drosophila* agar, 40g sucrose, 65g cornmeal, 25g Red Star whole yeast, 100 mL Karo lite corn syrup in 1200 mL dH_2_O) supplemented with 30% coconut oil as described previously [17] during the 4- 5-day aging period described above. For starvation experiments, male flies were food deprived on 1% agar for the last 24 hours of the aging period described above.

### Courtship analysis

Males and females were transferred to different sides of the divider in Aktogen^®^ courtship chambers by cooling them on ice and transferring with forceps. Once flies recovered from cooling and were walking normally in the chamber, the divider was removed allowing flies to interact. A video camera was placed over the chamber and all pairs were video recorded for three hours. Four chambers were always run together; two normal fed wild type male trials were always paired with two trials of one of the other three groups (food deprived, high fat diet, and *adp^60^*).

Videos were uploaded to a behavioral event recorder program, Observer^®^. The person scoring the behaviors did so blind to which group was being scored. The male behaviors analyzed were courtship latency, orientation, wing vibration, wing scissor, tapping, thrust, copulation attempt and copulation [19]. Orientation was defined as when the male oriented at any direction close to the female’s body and typically done for the entirety for courtship. The start of orientation signaled the start of courtship, and therefore the start of when the divider was removed allowing a male to see a female to when orientation began was considered the latency to courtship. Wing vibration was when the male’s wings vibrated horizontally and vertically at different angles. This behavior is known to create the “courtship song” which is important in the courtship process of *Drosophila* [21]. Wing scissor was when a fly’s wings were extended, crossed and re-crossed. Tapping was when the male touched any female body part, usually sideways, with its tarsus. Thrusting, which occurs during copulation and copulation attempts, was when the male grasped the female’s abdomen with his foretarsi, curled his abdomen downward and forward, making a fast contact with the female’s genitalia. Copulation occurred when a male stabilized on top of the female for a considerable amount of time, typically 10-15 minutes. A copulation attempt was counted when the male grasped the female’s abdomen and maintained his abdomen in contact with the female’s genitalia for more than one second (i.e. longer than a thrust), but did not complete the copulation (i.e. did not stabilize for an extended copulation period of time).

### Triglyceride, Glycogen and Protein Measurements

Triglyceride, glycogen and protein levels were measured as previously described [22]. Briefly, 2 male flies were homogenized in lysis buffer (140 mM NaCl, 50 mM Tris-HCl, pH 7.4, 0.1% Triton-X 100, and 1X protease inhibitor cocktail (Roche Life Sciences) by sonication. Thus, one “sample” consisted of the macromolecule levels measured in two flies with WT N = 16, *adp^60^* mutant N = 17, high fat diet N = 21, food deprived N = 17.

Triglycerides and protein were measured using the Triglycerides Liquicolor Test (Stanbio) and the BCA Protein Assay Kit (ThermoScientific), respectively, according to manufacturer’s instructions. Total glucose levels were measured using the Glucose Oxidase Reagent (Pointe Scientific) after treating samples with 8 mg/mL amyloglucosidase (Sigma) in 0.2M citrate buffer, pH 5.0 for 2 hours. Free glucose was measured in samples not treated with amyloglucosidase and glycogen levels were determined by subtracting free glucose from total glucose. Triglycerides and glycogen were normalized by dividing each by the total protein concentration of each sample.

### Statistical Analysis

All behaviors (except orientation latency) were corrected for the length of an individual’s courtship period (i.e. orientation period) and then compared statistically. The number of wing scissors, number of wing vibrations, and duration of copulation conformed to parametric assumptions and were analyzed via ANOVA. Total courtship time, courtship latency, wing scissor duration rate, wing vibration duration rate, and total number of thrusts were ln transformed to fit parametric assumptions and analyzed via ANOVA. An overall significant ANOVA test was followed by Tukey tests to identify which groups in the overall test were significantly different. Number of tapping behaviors, rate of tapping behavior, and rate of thrusting could not be transformed to fit parametric assumptions and so were analyzed via Kruskal Wallis. No Kruskal Wallis test was significant thus no multiple comparison testing was necessary for non-parametric analyses. Copulation frequency between groups was tested via a Pearson Chi-Square test followed up with a post hoc test using the residuals if the overall was found to be significant using a Bonferroni adjustment for multiple comparisons and thus an alpha level of 0.01. Energy data (glycogen, triglyceride and protein content) were analyzed via ANOVA as these data met all parametric assumptions. Significant overall effects were followed by Tukey multiple comparisons.

## Results

### Copulatory and Courtship Behavior

There was a difference in copulation frequency across groups (*X*^2^_3, N = 252_ = 42.395, P < 0.001; Fig 1). Post hoc tests revealed that the *adp^60^* mutants had a significantly lower frequency of copulations (13 / 75, 17.3%) relative to the other three groups (P = 0.001), while the high fat fed flies had a significantly greater proportion of copulations (24/33, 72.7%) than the other three groups (P < 0.001). Food deprived and wild type flies showed a similar proportion of copulations (food deprived: 20/30, 66.7%, P = 0.02; wild type: 62/114, 54.4%, P =0.03).

**Fig 1.**
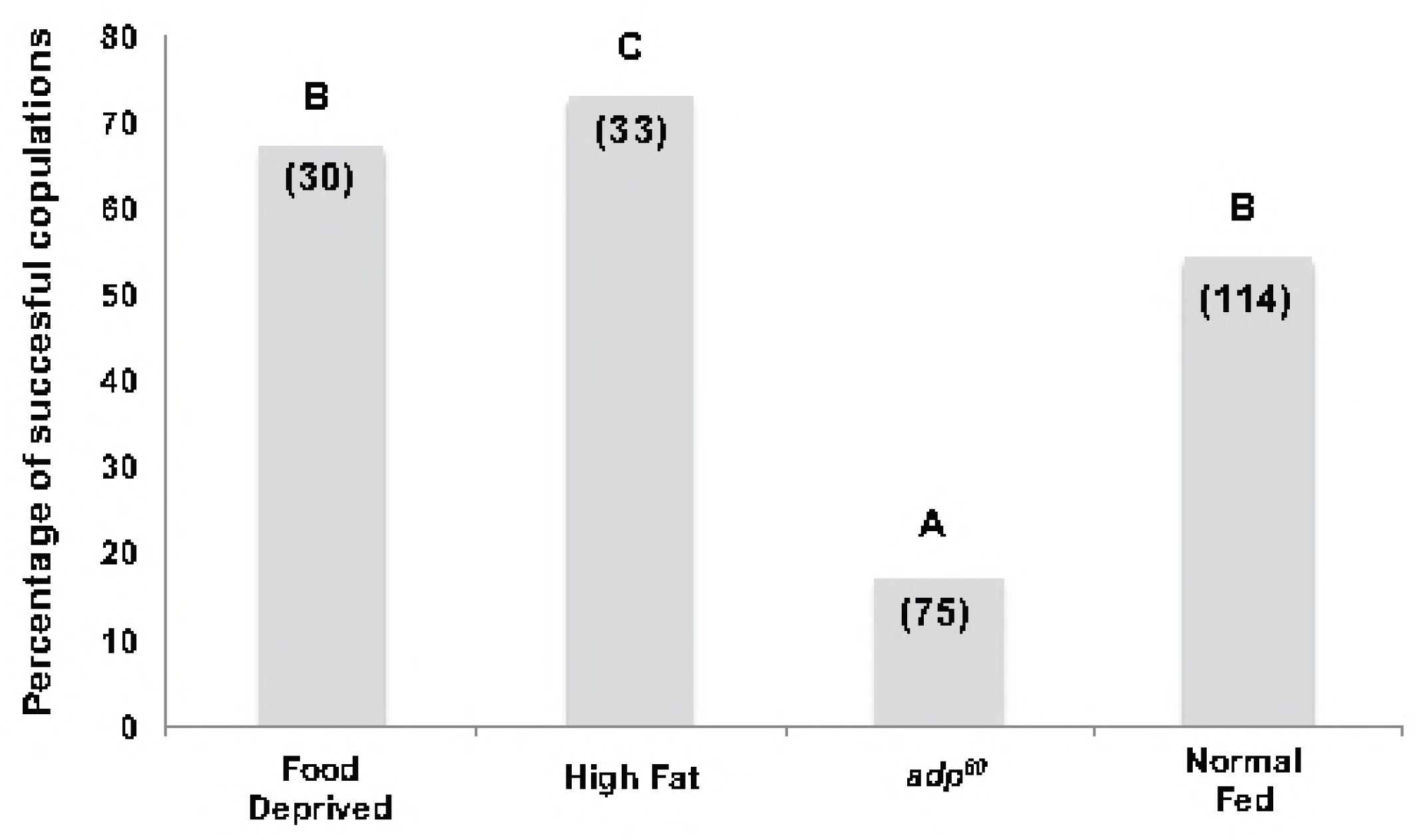
Percentage of successful copulations. This figure shows the percentage of successful copulations of food deprived, high fat fed, *adp^60^*, and normal fed flies.

Numbers in parentheses represent number of trials run in each group. Different letters indicate frequencies that were significantly different at P < .01 (based on Bonferroni adjustment; see text for details). Total courtship time among all four groups did not significantly differ (F _3,115_ = 0.789, P = 0.50). However, latency to court did significantly differ (F _3,115_ = 13.307, P < 0.001; Fig 2) with food deprived flies showing a significantly shorter latency time than either high fat fed flies (P = 0.036) or wild type flies (P < 0.001), but not relative to *adp^60^* mutants (P = 0.378). There were no significant differences in latency to court among high fat fed, wild type, and *adp^60^* mutant groups (P > 0.1 in all instances).

**Fig 2.**
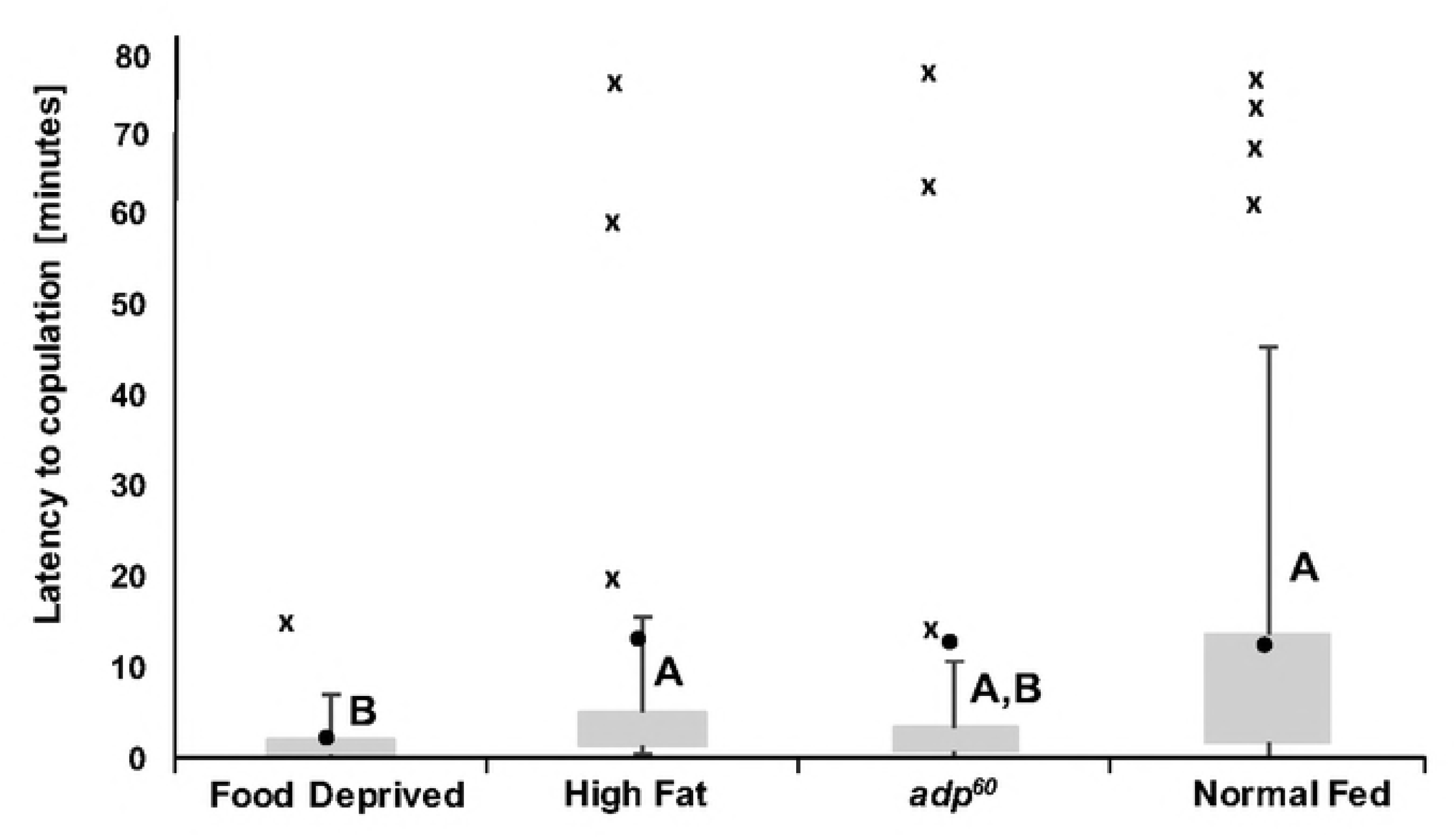
Courtship latencies. This figure shows the number of seconds required for flies in each group to engage in orientation behavior of food deprived, high fat fed, *adp^60^*, and normal fed flies. Boxes indicate interquartile ranges (IQR); whiskers indicate 1.5x IQR; “X” indicates points outside 1.5 ×; IQR; dark circles indicate mean. Different letters indicate groups that were significantly different at P < 0.05.

The total number of courtship behaviors did not significantly differ among all groups for any behavior (wing scissors: F _3,115_ = 0.522, P = 0.66; wing vibrations: F _3,115_ = 0.945, P = 0.42; thrusts: F _3,115_ = 0.212, P = 0.88; tapping: χ^2^ (3) = 0.25, P = 0.96), yet the rate these behaviors were conducted during courtship did significantly differ for some behaviors. Rate of wing scissoring significantly differed across groups (F _3,115_ = 10.711, P < 0.001; Fig 3) with the *adp^60^* mutant exhibiting a significantly lower rate than all other groups (vs. wild type: P = 0.002; vs. high fat flies: P < 0.001; and vs. food deprived flies: P< 0.001). Food deprived flies also showed a significantly higher rate of wing scissoring relative to wild type flies (P = 0.013), but was not different relative to high fat fed flies (P = 0.551). The rate of wing scissoring for wild types did not significantly differ from high fat fed flies (P = 0.357). Wing vibrations showed the same differences across groups as did wing scissoring. Specifically, the rate of wing vibrations significantly differed across groups (F _3,115_ = 12.343, P < 0.001) with the *adp^60^* mutant exhibiting a significantly lower rate than all other groups (P < 0.001 for all comparisons). Food deprived flies showed a significantly higher rate of wing vibration relative to wild type flies (P = 0.006), but was not different relative to high fat fed flies (P = 0.587). Wild type and high fat fed flies were again not different (P = 0.199). There was no significant effect on the rate of tapping (χ^2^(3) = 0.45, P = 0.92) or rate of thrusting (χ^2^ (3) = 4.503, P = 0.21) across all groups.

**Fig 3.**
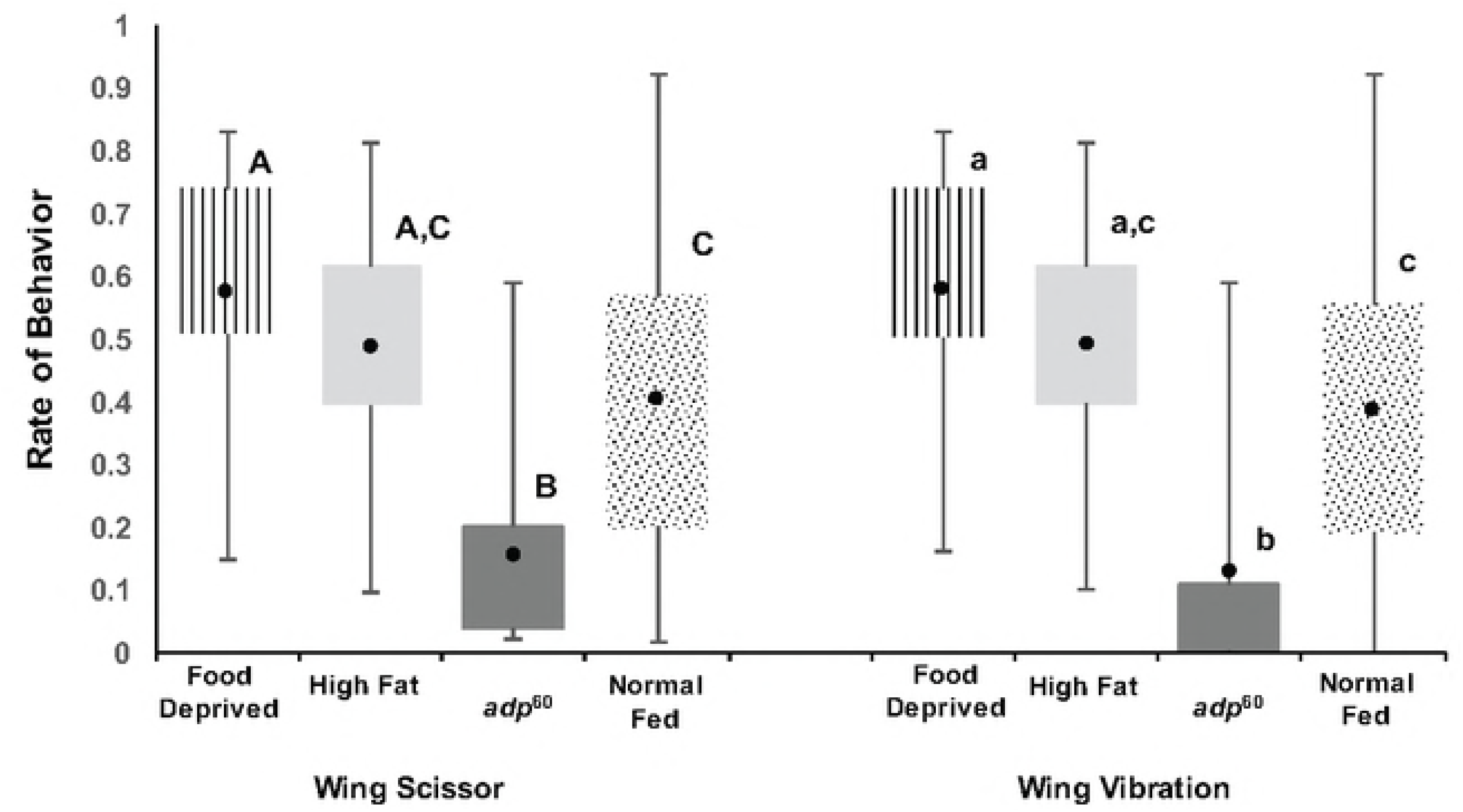
Courtship behavior rate. The rate at which courtship behavior (wing scissor and wing vibration) occurred for each group (vertical hatching = food deprived WT flies; light gray = high fat fed WT flies; dark gray = *adp^60^* mutant flies; stipple = normal WT fed). Rates are duration of behavior divided by duration of courtship (start of orientation to start of copulation). Boxes indicate IQR; whiskers indicate data range; dark circles indicate means. Different capital letters indicate groups that are significantly different for wing scissor behavior at P < 0.05. Different lower case letters indicate groups that are significantly different for wing vibration behavior at P < 0.05.

## Triglyceride and Glycogen levels

To confirm the efficacy of the high fat diet and the 24-hour food deprivation as well as the *adp^60^* mutation, triglyceride and glycogen levels were measured in these groups and compared to wild type flies fed normal food. Consistent with previous reports, there was a significant overall effect of treatment on the triglyceride per protein ratio (F _3,67_ = 173.293, P < 0.001; Fig 4A) with wild type flies showing a lower ratio than *adp^60^* mutants (P < 0.001) and high fat fed flies (P = 0.023), but a higher ratio compared to food deprived flies (P < 0.001) [16], [17], [23], [24]. Food deprived flies also showed a lower ratio compared to *adp^60^* mutants (P < 0.001) and high fat fed flies (P < 0.001). High fat fed flies showed a lower ratio compared to *adp^60^* mutants (P < 0.001). There was also a significant effect of fly/diet type on the glycogen per protein ratio (F _3,67_ = 38.524, P = 0.01; Fig 4B) with wild type flies showing a lower glycogen to protein ratio than *adp^60^* mutants (P < 0.001), a higher level than food deprived flies (P = 0.002) similar to that shown in [24] and no difference than the high fat diet fed group (P = 0.874). Food deprived flies also showed a lower glycogen to protein ratio compared to *adp^60^* mutants (P < 0.001) and high fat fed flies (P < 0.001). High fat fed flies showed a lower ratio compared to *adp^60^* flies (P < 0.001).

**Fig 4.**
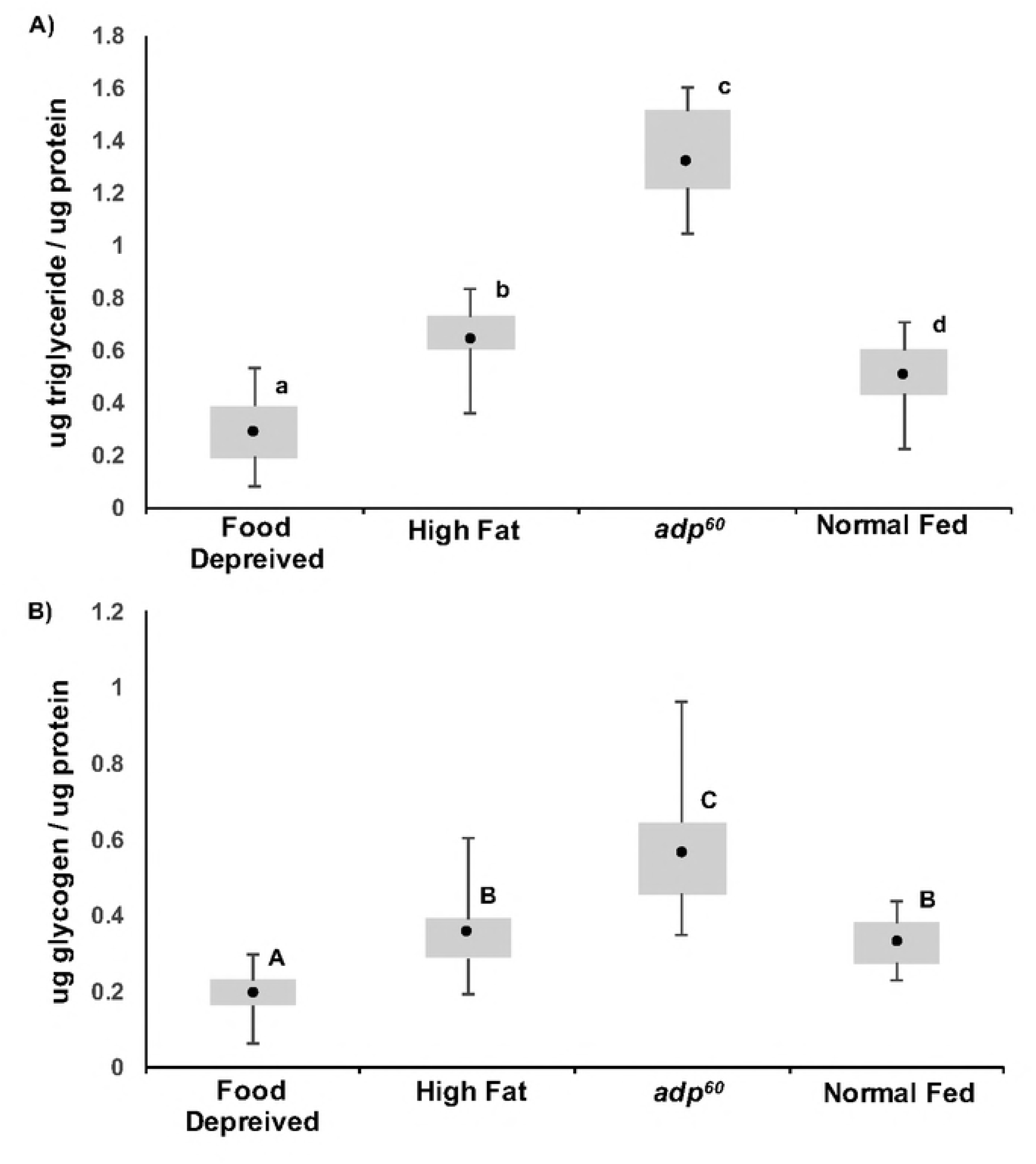
Fat and sugar per protein of flies. A) Triglyceride content per total body protein for food deprived, high fat fed, *adp^60^*, and normal fed flies. Different capital letters indicate significant differences at P < 0.05. B) Glycogen content per total body protein for food deprived, high fat fed, *adp^60^*, and normal fed flies. See Figure 2 for box and whisker details. Different lower case letters indicate significant differences at P < 0.05.

## Discussion

Availability of energy, both in terms of availability of nutrients and lipid and glycogen storage affected reproductive behaviors in *D. melanogaster* similarly. Contrary to our prediction, food deprived wild type flies had a decreased latency to court when compared to all other fly groups indicating that limiting caloric intake actually increases appetitive reproductive behavior. In addition, once courtship started, food deprived flies showed the greatest rate of courtship in terms of wing scissoring and wing vibrations which further supports this notion. However, this increased drive to reproduce did not result in more copulations. In fact, the high fat fed group enjoyed a significantly greater copulation frequency than all other groups, with the food deprived and normal fed flies showing similar copulation frequencies and *adp^60^* showing the least. Thus, while food deprived flies showed a greater drive and effort to reproduce, the higher copulation rate by high fat fed flies could indicate that flies fed diets high in fat are more attractive. This suggests that while energy availability influences male reproductive drive, other aspects of male appearance or courtship quality (two aspects known to influence female choice [25]) interact to impact reproductive success.

We suspect that the higher drive to reproduce shown by food deprived flies is due to the limited availability of metabolic resources, which could indicate a shortened time to live and thus a greater immediate investment in reproduction. Previous work has shown that food deprived flies and flies reared on highly nutrient restricted diets show decreased longevity compared to those raised on normal diets [26]. Flies who live shorter periods of time are also known to show increased drive to reproduce. For example, in *D. nigrospiracula*, male flies infected with parasites lived shorter lives but dedicated significantly more time to courting females than uninfected males [27]. Curiously, the courtship latency among normal fed wild type flies, high fat fed wild type flies and *adp^60^* mutants were all similar despite *adp^60^* mutants storing excess nutrients compared to wild type flies. However, courtship latency is indicative of courtship drive, and it is known that the lifespan of *adp^60^* homozygous males is similar to that of wild type males [28]. In addition, *adp^60^* mutants are starvation resistant [23]. Thus, *adp^60^* mutants do not appear to have as compromised reproductive potential based on survival as that of food deprived wild type flies.

Though longevity can be related to reproductive drive in species such as *Drosophila*, it remains unclear what is the direct cue to this increased drive – the actual intake of energy-rich molecules and/or the physiological availability of these metabolic reserves. Food deprived flies, who showed the shortest latency to court along with the highest rates of courtship behaviors (i.e. wing scissoring and wing vibrations), had the lowest food availability, and the lowest storage of both glycogen and triglycerides per body protein content. However, the *adp^60^* mutants, who had a constant availability of food but do not metabolize their triglycerides and glycogen in the same way as wild type flies (thus leading to an increased amount of each in our analysis) showed significantly less wing scissoring and wing vibrations. When considering these mutants show a similar reproductive drive to normal and high fat fed flies, potentially the availability of nutrients is involved with reproductive drive and courtship latency, while the appropriate storage and usage of these sources of energy once ingested is involved in carrying out courtship song caused by the wing vibration behavior. Perhaps combining diet alteration and the *adp^60^* mutation may allow the determination of whether the availability of nutrients or normal storage and usage of these nutrients is important for regulating courtship behaviors. However *adp^60^* mutants are known to have a “lethargic” phenotype where their overall mobility is less than that of wild type flies [23]. This phenotype may be due to this altered ability to metabolize nutrients and/or the gross obesity of these animals and could be related to the decreased wing scissoring and wing vibrations, but it did not affect the latency to court.

An important point to note is that this process could work differently in species with a different natural history. For example, it is well known that when physiological fuels are low in mammals, the reproduction cycle stops until such time as there is enough energy to put into reproduction [7]. This again is likely a function of longevity where such species can afford to wait for the proper resources. In mammals, it is well established that there are related, but independent mechanisms that link reproduction to food intake and physiological energy availability [6]. For example, kissipeptin, which is known to regulate GnRH and the reproductive axis, is influenced by ghrelin and PYY secretion (gastrointestinal hormones released upon ingestion) as well as by nutritional status [29]. Less is known about such pathways in *D. melanogaster*; however, there is some evidence to support that reproductive behavior is related to food availability (i.e. intake) and energy storage. For example, altering the amounts of protein and carbohydrates in the female diet will affect the rate of egg production in wild type *D. melanogaster* [30]. Schultzhaus [31] analyzed the effect of high fat diets on courtship in D. melanogaster and found that while feeding female flies a high fat diet had more negative effects on reproductive behavior than the same diet did on males, male judgment of female attractiveness was influenced by the high fat feeding. Considering only the wild type groups in the current study (food deprived, high fat fed, and normal fed), the nutrient storage data suggest that the reserves of glycogen, and not triglycerides, may serve as a cue for the change in sexual behavior we see in our study. Food deprived flies had lower glycogen and triglycerides than all other groups while normal fed and high fat fed flies courted similarly and had similar glycogen levels but different triglycerides. It follows that animals assess short term energy stores, such as glycogen, to establish levels of activity and not long term energy stores like triglycerides. We suspect that these limited metabolic resources culminated in a greater immediate investment in reproduction.

It is unclear why the *adp^60^* mutant contained the highest levels of glycogen and triglycerides yet courted the least, but many possibilities exist, each of which could contribute to and help explain the overall reduced mobility phenotype of the *adp^60^* mutant addressed above. One possibility is the physical structure of increased fat deposits decreases wing dexterity therefore affecting courtship song. Another is that the *adp^60^* mutation has additional unrealized effects on muscle activity. The *adp^60^* mutation results in chronic obesity with excess fat accumulation in both the larval and adult stages of development [16], [18], [23]; Figure 4A). Potentially these mutants do not mobilize these fuels in the same way as wild type flies as lipid metabolism in these mutants is suspected to be different throughout development (see [32] for further discussion). Any negative effect these metabolic pathways have on muscle function, in turn, could limit their ability to function properly and affect courtship. Lastly, when these male courtship songs are altered in this way, males might simply be choosing to decrease their song either because of a female’s lack of interest, or they are responding to their own song. While we cannot address which of these mechanisms is at play, they would all lead to reduced song and/or low quality song that could lead to an increased rejection by the female fly. In fact, the lack of courtship song from the *adp^60^* mutants could help explain the low copulation rates of this group via females “allowing” a copulatory grasp as various *Drosophila* species are known to reject males on the basis of song [33], [34].

While the lack of courtship song via wing vibrations could explain the significantly lower percentage of successful copulations for *adp^60^* mutants, food deprived flies did show the most wing vibrations, yet it was high fat fed flies who had the most copulations. Possibly females enable copulations based on multiple signals with courtship song being only one. Size of the male could be another (see [25] for further discussion], as well as number or quality of the cuticular hydrocarbons, which are known chemical communicators between males and females and can be affected by nutritional regime (reviewed in [35]). While high fat diets are known to not alter male cuticular hydrocarbons [31], the effect of *adp^60^* and food deprivation is not known. Larger males may generally indicate better quality sperm based on nutrition; however, Partridge et al.[25] also argued that larger males are more active and generally more persistent, so the increased copulation seen in the current study from the high fat fed group may be due to a form of male coercion.

## 1. Conclusions

Overall, we find that altering nutrient storage through diet or genetic manipulation has effects on courtship in the fruit fly, *D. melanogaster*. The increased drive to reproduce by food deprived flies was unexpected, but is reasonable based on species’ reproductive investment and longevity. We suspect that such findings would be different in species with a different natural history, such as those with a greater longevity where one can afford to wait for the proper resources to become available. More research is needed to identify the exact physiological cue that activates reproductive behavior across species with such varied life histories to better understand reproductive motivation in the context of natural history. We believe utilizing a combination of manipulation of animal diets and the use of mutants to manipulate physiological pathways can help illuminate the resource mechanisms that drive the reproduction axis. Species such as *D. melanogaster* are particularly powerful in this regard as such mutants exist both naturally, such as the *adp^60^* mutants used here, and can be artificially created to probe the effects of specific genes and metabolic processes on organismal reproduction.

## 2. Acknowledgements

We would like to thank Grzegorz Polak and Justin Palermo for help with fly husbandry. This research did not receive any specific grant from funding agencies in the public, commercial, or not-for-profit sectors.

## References

1. Kohsaka A, Laposky AD, Ramsey KM, Estrada C, Joshu C, Kobayashi Y, et al. High-fat diet disrupts behavioral and molecular circadian rhythms in mice. Cell Metab. 2007;6:414–421. DOI: 10.1016/j.cmet.2007.09.00.

2. Lim RS, Eyjólfsdóttir E, Shin E, Perona P, Anderson DJ. How food controls aggression in Drosophila. PLoS One 2014;9:e105626. DOI:10.1371/journal.pone.0105626

3. Redman LM, Heilbronn LK, Martin CK, De Jonge L, Williamson DA, Delany JP, et al. Metabolic and behavioral compensations in response to caloric restriction: implications for the maintenance of weight loss. PLoS One 2009;4:e4377. DOI: 10.1371/journal.pone.000437.

4. Bronson F. Puberty and energy reserves: a walk on the wild side. In: Wallen, K., Schneider JE, editors. Reproduction in Context: Social and Environmental Influences on Reproduction. Cambridge, Massachusetts; London, England: MIT Press; 2000. p.15–33.

5. Gittleman JL, Thompson SD. Energy allocation in mammalian reproduction. Am. Zool. 1988;28:863–875. DOI: 10.1093/icb/28.3.86.

6. Schneider JE. Energy balance and reproduction. Physiol. Behav. 2004; 81:289–317. DOI: 10.1016/j.physbeh.2004.02.00.

7. Schneider JE, Wade GN. Inhibition of reproduction in service of energy budget. In: Wallen, K., Schneider JE, editors. Reproduction in Context: Social and Environmental Influences on Reproduction. Cambridge, Massachusetts; London, England: MIT Press; 2000 p.15–33.

8. Martin TE. Food as a limit on breeding birds: a life-history perspective. Annu. Rev. Ecol. Syst. 1987;18:453–487.

9. Sibly RM, Witt CC, Wright NA, Venditti C, Jetz W, Brown JH. Energetics, lifestyle, and reproduction in birds. Proc. Natl. Acad. Sci. 2012;109:10937–10941. DOI: 10.1073/pnas.120651210.

10. Lind CM, Beaupre SJ. Male Snakes Allocate Time and Energy according to Individual Energetic Status: Body Condition, Steroid Hormones, and Reproductive Behavior in Timber Rattlesnakes, Crotalus horridus. Physiol. Biochem. Zool. 2015;88:624–633. DOI: 10.1086/68305.

11. Van Dyke JU, Griffith OW, Thompson MB. High Food Abundance Permits the Evolution of Placentotrophy: Evidence from a Placental Lizard, Pseudemoia entrecasteauxii. Am. Nat. 2014;184:198–210. DOI: 10.1086/67713.

12. Ottinger MA, Mobarak M, Abdelnabi M, Roth G, Proudman J, Ingram DK. Effects of calorie restriction on reproductive and adrenal systems in Japanese quail: are responses similar to mammals, particularly primates? Mech. Ageing Dev. 2005; 126:967–975. DOI: 10.1016/j.mad.2005.03.01.

13. Cheng D, Chen L, Yi C, Liang G, Xu Y. Association between changes in reproductive activity and D-glucose metabolism in the tephritid fruit fly, Bactrocera dorsalis (Hendel). Sci. Rep. 2014; 4(7489):1–9.14. DOI: 10.1038/srep0748.

14. Warburg MS, Yuval B. Effects of energetic reserves on behavioral patterns of Mediterranean fruit flies (Diptera: Tephritidae). Oecologia 1997;112:314–319. DOI: 10.1007/s00442005031.

15. Baker KD, Thummel CS. Diabetic larvae and obese flies—emerging studies of metabolism in Drosophila. Cell Metab. 2007;6:257–266.

16. Häder T, Müller S, Aguilera M, Eulenberg KG, Steuernagel A, Ciossek T, et al. Control of triglyceride storage by a WD40/TPR-domain protein. EMBO Rep. 2003;4: 511–516. DOI: 10.1038/sj.embor.embor83.

17. Birse RT, Choi J, Reardon K, Rodriguez J, Graham S, Diop S, et al. High-fat-diet-induced obesity and heart dysfunction are regulated by the TOR pathway in Drosophila. Cell Metab. 2010;12:533–544. DOI: 10.1016/j.cmet.2010.09.01.

18. Reis T, Van Glist MR, Hariharan IK. A buoyancy-based screen of Drosophila larvae for fat-storage mutants reveals a role for Sir2 in coupling fat storage to nutrient availability. PLoS Genet. 2010; 6(11): e1001206. DOI: 10.1371/journal.pgen.1001206.

19. Cobb M, Connolly K, Burnet B. Courtship behaviour in the melanogaster species sub-group of Drosophila. Behaviour. 1985; 95:203–230. DOI: 10.1163/156853985X0013.

20. Kauffman, A.S., Rissman, E.F., 2004. A critical role for the evolutionarily conserved gonadotropin-releasing hormone II: mediation of energy status and female sexual behavior. Endocrinology 145, 3639–3646. DOI: 10.1210/en.2004-014.

21. Ewing AW, Bennet-Clark H. The courtship songs of Drosophila. Behaviour 1968; 31: 288–301. DOI: 10.1163/156853968X0029.

22. Gingras RM, Warren ME, Nagengast AA, DiAngelo JR. The control of lipid metabolism by mRNA splicing in Drosophila. Biochem. Biophys. Res. Commun. 2014;443: 672–676. DOI: 10.1016/j.bbrc.2013.12.02.

23. Suh JM, Zeve D, McKay R, Seo J, Salo Z, Li R, et al. Adipose is a conserved dosage-sensitive antiobesity gene. Cell Metab. 2007;6:195–207.

24. Schwasinger-Schmidt TE, Kachman SD, Harshman LG. Evolution of starvation resistance in Drosophila melanogaster: measurement of direct and correlated responses to artificial selection. J. Evol. Biol. 2012;25:378–387. DOI: 10.1111/j.1420-9101.2011.02428.

25. Partridge L, Ewing A, Chandler A. Male size and mating success in Drosophila melanogaster: the roles of male and female behaviour. Anim. Behav. 1987;35: 555–562. DOI: 10.1016/S0003-3472(87)80281-.

26. Magwere T, Chapman T, Partridge L. Sex differences in the effect of dietary restriction on life span and mortality rates in female and male Drosophila melanogaster. J. Gerontol. A. Biol. Sci. Med. Sci. 2004;59:B3–B9. DOI: 10.1093/gerona/59.1.B.

27. Polak M, Starmer WT. Parasite–induced risk of mortality elevates reproductive effort in male Drosophila. Proc. R. Soc. Lond. B Biol. Sci. 1998;265:2197–2201. DOI: 10.1098/rspb.1998.055.

28. Doane WW. Developmental physiology of the mutant female sterile (2) adipose of Drosophila melanogaster. I. Adult morphology, longevity, egg production, and egg lethality. J. Exp. Zool. Part Ecol. Genet. Physiol. 1960;145:1–21.DOI: 10.1002/jez.140145010.

29. Fernandez-Fernandez R, Martini A, Navarro V, Castellano J, Dieguez C, Aguilar E, et al. Novel signals for the integration of energy balance and reproduction. Mol. Cell. Endocrinol. 2006; 254: 127–132.

30. Lee KP, Simpson SJ, Clissold FJ, Brooks R, Ballard JWO, Taylor PW, et al. Lifespan and reproduction in Drosophila: new insights from nutritional geometry. Proc. Natl. Acad. Sci. 2008;105:2498–2503. DOI: 10.1073/pnas.071078710.

31. Schultzhaus JN, Bennett CJ, Iftikhar H, Yew JY, Mallett J, Carney GE. High fat diet alters Drosophilia melanogaster sexual behavior and traits: decreased attractiveness and changes in pheromone profiles. Sci. Rep. 2018; 8:5387. DOI:10.1038/s41598-018-23662-2

32. Teague BD, Clark AG, Doane WW. Developmental analysis of lipids from wild-type and adipose60 mutants of Drosophila melanogaster. J. Exp. Zool. Part Ecol. Genet. Physiol. 1986;240:95–104.

33. Hoikkala A, Aspi J, Suvanto L. Male courtship song frequency as an indicator of male genetic quality in an insect species, Drosophila montana. Proc. R. Soc. Lond. B Biol. Sci. 1998;265:503–508. DOI: 10.1098/rspb.1998.032.

34. Ritchie MG, Saarikettu M, Livingstone S, Hoikkala A. Characterization of female preference functions for Drosophila montana courtship song and a test of the temperature coupling hypothesis. Evolution 2001;55:721–727. DOI: 10.1554/0014-3820(2001)055[0721:COFPFF]2.0.CO;.

35. Ferveur JF. Cuticular hydrocarbons: their evolution and roles in Drosophila pheromonal communication. Behav. Genet. 2005;35:279–295. DOI: 10.1007/s10519-005-3220-.

